# AutoMolDesigner for Antibiotic Discovery: An AI-based Open-source Software for Automated Design of Small-molecule Antibiotics

**DOI:** 10.1101/2023.09.27.559854

**Authors:** Tao Shen, Jiale Guo, Zunsheng Han, Gao Zhang, Qingxin Liu, Xinxin Si, Dongmei Wang, Song Wu, Jie Xia

## Abstract

Discovery of small-molecule antibiotics with novel chemotypes serves as one of the essential strategies to address antibiotic resistance. Although a considerable number of computational tools committed to molecular design have been reported, there is a deficit in the holistic and efficient tool specifically developed for small-molecule antibiotic discovery. To address this issue, we report AutoMolDesigner, a computational modeling software dedicated to small-molecule antibiotic design. It is a generalized framework comprising two functional modules, i.e., generative deep learning-enabled molecular generation and automated machine learning based-antibacterial activity/property prediction, wherein individually trained models and curated datasets are out-of-the-box for whole cell-based antibiotic screening and design. It is open-source thus allows for the incorporation of new features for flexible use. Unlike most software programs based on Linux and command lines, this application equipped with Qt-based graphical user interface can be run on personal computers with multiple operating systems, making it much easier to use for experimental scientists. The software and related materials are freely available at GitHub (https://github.com/taoshen99/AutoMolDesigner) and Zenodo (https://zenodo.org/record/8366085).

## INTRODUCTION

The evolutionary multidrug-resistant (MDR) bacteria have become an emerging threat to public health^1^. Among these bacteria, the ESKAPE (*Enterococcus faecium*, *Staphylococcus aureus*, *Klebsiella pneumoniae*, *Acinetobacter baumannii*, *Pseudomonas aeruginosa*, and *Enterobacter*) pathogens^2^ account for many fatal infections. Known chemotypes of antibiotics rapidly lose efficacy to these pathogens due to the developed resistance mechanisms through natural selection^3^. Accordingly, novel chemotypes of antibiotics that are widely acknowledged as effective to overcome antibiotic resistance are at an urgent need. In history, novel-chemotype antibiotics are mainly discovered by three strategies. The first strategy dates back to half a century ago, the exploitation of natural products (secondary metabolites) isolated from environmental microbes contributed to most of the small-molecule antibiotics in clinic^4^. However, this strategy is becoming insufficient now since the same chemotype (skeleton) is repetitively discovered and structural modification does not lead to better antibiotics than the natural product itself^5^. With the emerging of combinatorial chemistry at the beginning of the 21st century, the second strategy was target assay-based high-throughput screening (HTS)^1^ of large-scale chemical library. The target-based approach achieved notable success in drug research and development for other diseases. Unfortunately, this strategy seems not fruitful in antibiotic discovery, as only three new chemotypes of antibiotics received approval^6–8^ in the last 20 years and two of them are still derived from natural products. The third strategy is phenotype-based antibiotic screening, in particular the whole cell-based antibiotic screening, i.e. directly screening chemical libraries for bioactive compounds against the whole bacteria instead of a protein target, which holds the promise to circumvent the pitfall of target assay-based screenings that do not consider antibiotic-likeness^9^.

Machine learning (ML), as the core of AI, is comprised of three basic subfields including supervised learning, unsupervised learning and reinforcement learning (RL). Correspondingly, the application of ML to whole cell-based antibiotic discovery can be divided into two aspects according to the applied algorithms^10^. Quantitative structure–activity relationships (QSAR) modeling (also called molecular bioactivity prediction) is the most important supervised learning task in this field^11^. It utilizes whole-cell activity prediction models to prioritize compounds from chemical libraries. A few cases in antibacterial hit identification have been reported. For example, Wang *et al.* utilized molecular descriptors and fingerprints to characterize known 2,066 compounds with activity against methicillin-resistant *S. aureus*, and they built models with four classical ML algorithms including naive Bayesian, support vector machine, recursive partitioning, and k-nearest neighbors^12^. The best models were eventually applied to virtual screening of a chemical library comprised of about 7,500 compounds, which contributed to several antibacterial hits. Recently, molecular property prediction based on deep learning-enabled automated featurization was reported^13^. Chemprop is developed to extract bond-level topological information from graph representation based on the algorithm of directed-message passing neural network (D-MPNN)^14^. With two kinds of cell-based antibiotic datasets compiled for model training, Chemprop was respectively applied to virtual screening campaigns for new antibiotics. As a result, the research teams have discovered two novel hits which are either broad-spectrum antibiotics^15^ or narrow-spectrum antibiotics against *A. baumannii* with a new mode of action^16^. With the renaissance of artificial intelligence (AI), *de novo* design of antibacterial molecules becomes an emerging field, wherein self-supervised learning, a kind of unsupervised learning, has been leveraged for molecular generation. Currently, most research is focused on *de novo* design of antimicrobial peptides and those applied strategies are pretty effective^17^. As few cases of success were reported, deep learning-based design of small-molecule antibiotics seems quite challenging. Segler *et al.* pioneered the framework of recurrent neural network (RNN)-based chemical language models (CLMs) for small-molecule generation^18^. In their study, the *in silico* design of novel small-molecules antibiotics against *S. aureus*. was used to illustrate the “Design-Synthesis-Test Cycle”. The ML-based prediction model and the transfer learning strategy were coupled to generate a focused chemical library including 60,988 virtual molecules with potential antibacterial activity. This important work provided an early insight into the small-molecule antibiotic design through generative deep learning. However, their ML-based prediction model covered antibacterial activity only and neglected antibiotic-likeness, thus the designed molecules may not be ideal. In terms of public availability, no source code or application was provided. Recently, Sowmya *et al.* developed a semi-supervised deep molecular generation model to explore structure-based molecular design for *Mycobacterium tuberculosis*. A key ML model for drug-target affinity prediction was used to reward the generative process through RL^19^. They have shown that more than 75% generated molecules are highly analogous to known inhibitors and contained the essential pharmacophore features that would interact with the binding site residues of chorismate mutase. Unfortunately, they did not take the whole-cell antibacterial activity into consideration but only emphasized target binding of the designed molecules. Therefore, it is not convincing that the designed molecules by the model are active for the bacteria. Similar to the work of Segler *et al*., their model was not made publicly available as well.

To the best of our knowledge, the current development of AI-based computational tools for antibiotic design and screening is far from satisfaction. Two main issues are as follows: (1) All the whole cell-based models in previous study are mainly focused on antibacterial activity profiling while lack the consideration of antibiotic-likeness^20^, a significantly important aspect of antibiotics. This aspect is also overlooked in molecular generation-based small-molecule antibiotic design from a target structure-centered perspective. (2) The computational tools and models for antibiotic design and screening are not easy to access by users. Most of them required programming skills and high-performance server to deploy for application. Recently, some web servers such as AIScaffold^21^ and Chemistry42^22^ were developed for molecular design. Basically, they are applicable to the building of antibiotic-focused AI models, but they all require commercial licenses.

To address these issues, we developed AutoMolDesigner for small-molecule antibiotic design and screening, which consisted of two functional modules, i.e., a remastered RNN aiming at focused library generation for small-molecule antibiotics and an automated machine learning (AutoML) framework named AutoGluon^23^ for molecular property prediction. Both classification and regression models are out-of-the-box for accurate prediction of antibacterial activity and antibiotic-likeness, with cytotoxicity and plasma protein binding (PPB) as representatives. To make the models easy to use, we incorporated the modules including data curation, model evaluation and molecular visualization into AutoMolDesigner. In addition, we designed an intuitive graphical user interface (GUI) for both Windows and MacOS platform, along with the corresponding command-line interface (CLI). This software was also tested on personal computers with various device configurations. It can be run on a laptop without independent graphics processing unit (GPU) and achieve close performance to that on high-performance servers.

## RESULTS AND DISCUSSION

Three main features of AutoMolDesigner are summarized below:

1. A generalized framework was implemented to realize automated molecular design through focused library generation and molecular property prediction;
2. Specific datasets and models for antibiotic discovery were established for the out-of-box use.
3. Software operating logics were simplified, intuitive GUI was provided and fluent user experience on personal computer was guaranteed.

Figure 1 depicts the workflow of this software, and corresponding interfaces of two main sub-tools are displayed in Figure S1. Herein, we take the automated design of antibiotic-like small molecules with activity against *Escherichia coli* (*E. coli.*) as a case to demonstrate the usage of the two tools.

**Figure 1.**
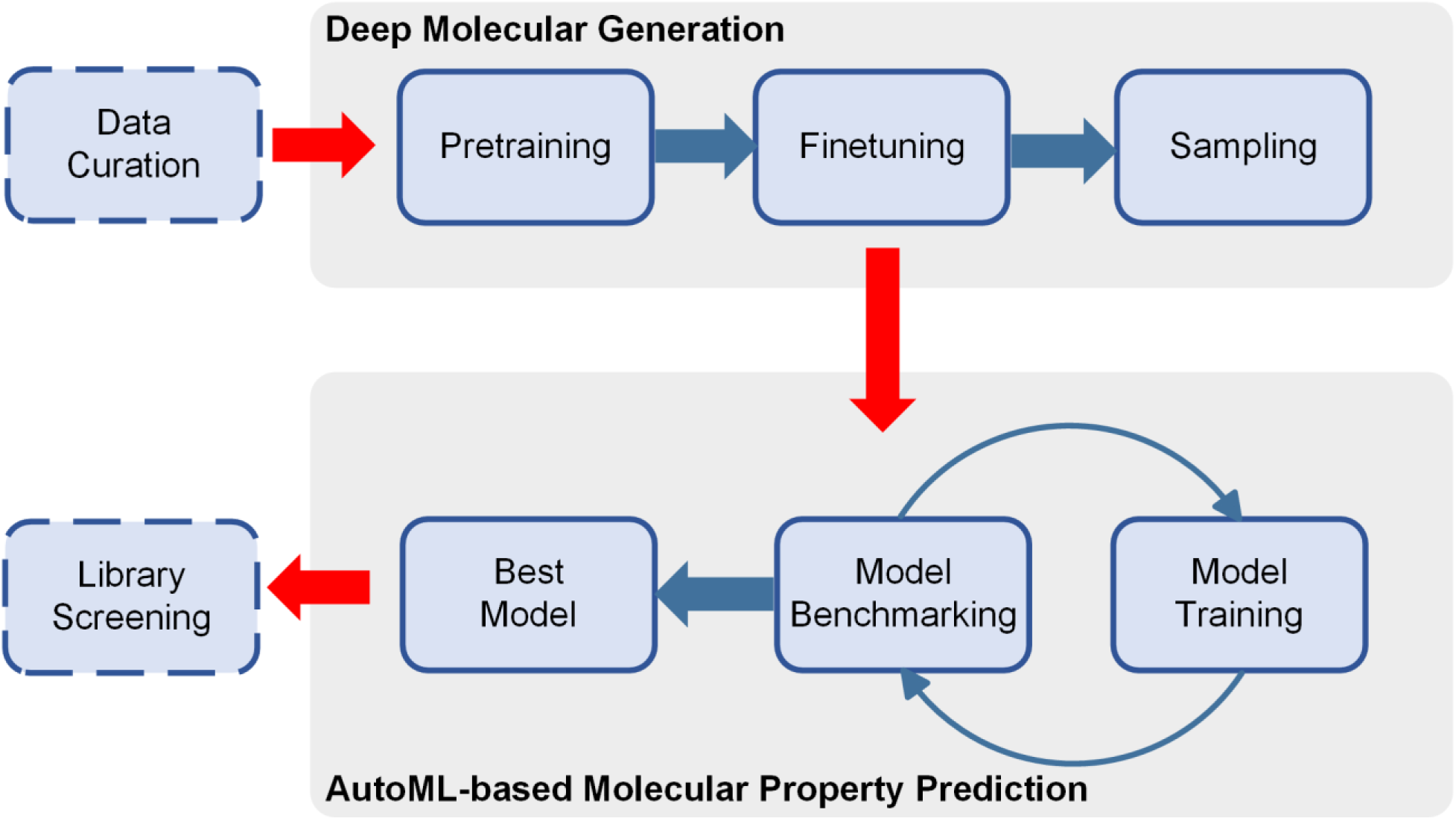
Workflow of automated molecular design implemented in AutoMolDesigner.

### Tool I: Generating Focused Library for Small-molecule Antibiotics

#### Data Curation

Before model training, data curation implemented in the first tab named “Data Preparation” should be conducted first to provide accurate and uniform data for ML model. Apart from the default curation such as de-duplication, salt removal, and molecular standardization, the other two custom functionalities are provided as well. First, neutralization can be operated on molecules that are used for molecular generation. Ionization states at a certain pH range can be enumerated by Dimorphite-DL^24^, an open-source Python package for drug-like molecular ionization. This functionality is indispensable for molecular property prediction. Second, molecular data augmentation is available, which performs the simplified molecular input line entry system (SMILES) enumeration. It has been proved effective to enhance model performance for both supervised and unsupervised tasks, especially for molecular property prediction^25^ and transfer learning in low data regimes^26^.

We retrieved a large amount of bioactive data from ChEMBL database^27^ (version 32, https://www.ebi.ac.uk/chembl/, accessed March 2023) for model pretraining. The default data curations, molecular neutralization and augmentation by ten folds were conducted to give 7,230,290 data points (“Dataset for Pretraining”). In terms of data used for model finetuning (also named transfer learning), we retrieved active molecules whose measured minimum inhibitory concentration (MIC) values against *E. coli.* were no greater than 32 μg/mL from ChEMBL database (version 32, accessed March 2023). Apart from the same curation as that for the pretraining data, we additionally removed the molecules whose molecular weight is greater than 600 and number of rotatable bonds is more than 20 by referring to the compilation of MUBD-HDACs^28^. A total of 13,243 data points were obtained and their distribution can be visualized from Figure S2. The MIC value of 1 μg/mL (-log_2_^MIC^ value of 0) was used as the cutoff to discriminate between the compounds with “moderate activity” and those with “high activity”. The molecules labeled as “high activity” were later used to train the molecular property prediction model. The molecules labeled as “moderate activity” accounted for about two thirds of all compounds. The structural clustering of all the 13,243 molecules based on Morgan2 (Extended-Connectivity Fingerprints with radius 2, ECFP_4) fingerprints^29^ was carried out, resulting in a total of 570 clusters. Table S2 lists the cluster attribute of each molecule. Moreover, a set of molecules constituted by the compounds with the lowest MIC values from 6 largest clusters along with the compounds from 6 smallest clusters (size = 1) are displayed in Figure S3. As expected, 6 compounds from those large clusters all belong to well-studied chemotypes of small-molecule antibiotics including quinolone, β-lactam, thiazolidinone, propargyl alcohol and diacetylene. In comparison, the other 6 compounds from the smaller clusters have novel scaffolds but show less potency. The above data indicate the dataset is diverse. It not only covers the well-known chemotypes, but also includes rare but structurally novel antibiotics. It should be noted that only the active molecule with the lowest MIC value from each cluster was used for transfer learning. Thus, the diverse subset (“Dataset for Finetuning”) was composed of 570 compounds.

#### Molecular Generation

By referring to several recently reported prospective campaigns that made full use of deep molecular generation^30, 31^, we designed the main interface named molecular generation (MG) based on an autoregressive model that took a remastered RNN as its core network. Three sub-modules that executed model pretraining, model finetuning and molecular sampling constituted the whole workflow of molecular generation, and they were implemented in three independent tabs, i.e., “MG Pretrain”, “MG Finetune” and “MG Sample”.

Here, we demonstrate the molecular generation beginning with “MG Pretrain”. This submodule aims at learning generalized features of drug-like molecules to enable the unbiased production of stochastic molecules with possible bioactivity. Despite multiple hyperparameter options are available at this tab, only advanced users are recommended to adjust the architecture. Users who focus on practice are suggested to use the generalized model parameters that we have provided at GitHub. The model was pretrained with the “Dataset for Pretraining”, and its architecture is described in the section of “Materials and Methods, Deep Molecular Generation” in Supporting Information. To follow the routine of the scientific community, we performed benchmarking of our MG model with MOSES^32^. Table S1 shows that our remastered RNN was superior to CharRNN but fell behind MolGPT^33^ in molecular validity. It generated more novel molecules and achieved lower Frechet ChemNet Distance (FCD) score compared with MolGPT. Lower value of FCD score indicates the better capability of model in learning dataset statistics. Notably, MolGPT contained far more network parameters than RNN as its basic architecture was generative-pretraining Transformers, which is computationally expensive and basically trained on high-performance servers. Our model performance was close with these two models in terms of other metrics, indicating the robustness of our remastered MG model implementation.

Following the model pretraining, it came to the transfer learning that plays a key role in practice of molecular generation. Deep learning model attempts to learn from the user-customized data that has shared features (for example, with bioactivity against a specific target, protein family or even phenotype). Ideally, the direction of molecular generation will be steered towards the desired chemical space. It is worth noting that number of iteration (epoch) could be customized in this tab but early stopping technique was also provided to prevent overfitting, thus model training may stop before the preset epoch is met. Considering that model finetuning is the most demanding task on computing capacity in practice, we compared runtime performance on various device configurations using exactly the same parameters and finetuning dataset (“Dataset for Finetuning”). Table S3 shows that it took only 222 seconds to complete this task on a Windows laptop with a common central processing unit (CPU), and 35 seconds more time on an Apple computer, indicating the acceptable running speed of our software. The high-performance server only ran for 7 seconds to complete the task, indicating the introduction of GPU could lead to obvious improvement. The parameters of the finetuned model are also available at GitHub.

At last, “MG Sample” submodule was developed to sample new molecules from the pretrained or finetuned model. The “Temperature Sampling” function is borrowed from other CLMs to improve the sampling process^31^. More molecules with unusual atom types and novel scaffolds will be generated at lower temperature while invalid molecules may also be generated more frequently. Since not every SMILES sampled from the trained models can be converted to valid molecules, the “MG Post-process” module is provided here to measure quality of the generated molecules and perform data curation. The metrics^32^ including “Validity” (fraction of valid molecules in all generated molecules) and “Uniqueness” (fraction of non-duplicated molecules in all valid molecules) will be shown in a message box after the removal of invalid SMILES and the duplicates. The metric of “Novelty” will also be shown by calculating the number of unique SMILES after de-duplication from the “Dateset for Finetuning” (Figure S4). By default, the “MG Sample” submodule sampled 30,000 molecules by using the trained model.

#### Library Analysis

To analyze the focused library generated for antibiotics against *E. coli.*, we used the “MG Sample” submodule to sample 1,000 molecules and then used uniform manifold approximation and projection^34^ (UMAP) to visualize the chemical space characterized by Functional-Class Fingerprints with radius 3 (FCFP_6). The chemical space covered 967 valid molecules sampled from the pretrained model, 910 valid and de-duplicated molecules sampled from the finetuned model and 570 molecules from “Dataset for Finetuning”. Figure S5 shows that most molecules sampled from the finetuned model were adjacent to those known antibiotics in chemical space, proving that the model was able to capture the antibiotic-specific features through transfer learning. Notably, a few molecules generated by finetuning occupied the space of the molecules generated by pretraining but were not that far from the space of known antibiotics, thus promising small-molecule antibiotics with novel chemotypes may be discovered from these molecules.

### Tool II: AutoML-based Molecular Property Prediction for Small-molecule Antibiotics

Although Tool I could generate antibiotic-like molecules as a focused library, it is still necessary to narrow down the chemical space with ML-based molecular property prediction. Herein, we applied AutoML for our purpose^35^. Such a discipline aims at building task-oriented ML models and further maximizing overall performance with ensemble learning methods including bagging, boosting and stacking. Its main advantages include minimal human intervention, highly parallel model training and optimization as well as universal applicability. Among the AutoML frameworks, AutoGluon is an open-source application developed and distributed by Amazon Co. Ltd., which has achieved state-of-the-art performance on the tabular data benchmarks^23^. More importantly, this tool was recently proved applicable to predict pharmacokinetics-related molecular properties^36^. Accordingly, AutoGluon combined with molecular descriptors/fingerprints was used here to build antibiotic-related molecular property prediction models. As shown in Figure S1B and S6, the “AutoGluon” sub-interface contains two modules, i.e., “Model Training” and “Model Prediction”.

#### Model Training

The “Model Training” module provides five kinds of molecular descriptors/fingerprints, i.e., RDKit 2D^14^, normalized RDKit 2D, ECFP_4, FCFP_6 and MACCS structural keys. The other adjustable arguments are derived from AutoGluon. The “Model Quality Preset” argument determines model performance controlling number of trained models and hyperparameter configurations. This “Model Quality Preset” has four levels ranging from low level to high level, i.e., “medium_quality”, “good_quality”, “high_quality” and “best_quality”. More computational resources are required if the level is higher. “Evaluation Metric” argument defines the metrics for model training, i.e., F1 score and area under receiver operation characteristic curve (AUROC) for classification and mean absolute error (MAE) and root mean squared error (RMSE) for regression. The “Time Limit” argument sets the maximum amount of time for model training. The training will be forced to stop if the elapsed time is beyond the limit. The “Deployment optimization” argument is used to reduce computational cost by the removal of redundant models but may slightly reduce model performance.

Tabular data is accepted for model training. Its first column is headed with “SMILES” containing molecules represented by SMILES. The second column is headed with “label” containing either 0/1 for binary classification or continuous values for regression. Raw data should be firstly prepared by “Data preparation” module, where Comma-Separated Values (*.csv) and Microsoft Excel (*.xlsx) are two supported formats for input. By default, the whole data are used for training while random splitting of the datasets into a training set and a test set is also available. All the trained ML models are saved under the directory named “ag_models”.

#### Model Prediction

There are two modes in “Model Prediction” module. For retrospective prediction, users can provide tabular data in the same format as the training data. The model performance is measured by confusion matrix, accuracy, AUROC, Matthews correlation coefficient (MCC) and F1 score for classification and MAE, RMSE, R square (*R*^2^) and median absolute error (MedianAE). For prospective prediction, a plain text file containing molecules represented by SMILES (*.smi, one sequence per row) is required. Both a csv file and a Structure Data File (*.sdf) are used to save the predicted values. To demonstrate the functionality of the AutoGluon-based ML tools, three cases for predicting antibacterial activity, cytotoxicity and PPB are discusses below.

#### Antibacterial Activity

The aforementioned studies have proven the feasibility of discovering new antibiotics using bacterial phenotype-based ML models. In this case, 4,774 known antibiotics with “high activity” (MIC: less than 1 μg/mL) against *E. coli.* were used for modeling. They were firstly used to make benchmarking sets by MUBD-DecoyMaker2.0^37^, a publicly accessible GUI application focused on making trustworthy datasets for virtual screening. The benchmark for model training was comprised of 501 positive data points (unbiased ligands) and 19,539 negative data points (unbiased decoys), which simulated the virtual screening scenario in real world (i.e., a low hit rate in HTS, Table S4). Since Chemprop is an established ML tool for antibacterial activity prediction^14^, we carried out comparative studies between this tool and AutoGluon. Table S5 shows that the best performance was achieved by Morgan2-based AutoGluon model, with the AUC of 0.937 and MCC of 0.687, which significantly outperformed the best Chemprop model, with the AUC of 0.877 and MCC of 0.552. Notably, the best Chemprop model was enhanced by ensemble learning and molecular-level feature^14^ (normalized RDKit 2D). In comparison, we set the “Time limit” argument of the best AutoGluon model to 3,600 seconds, indicating that the model performance could be further improved if extra time is allocated. In terms of runtime, it took 27 seconds to train RDKit 2D-based AutoGluon model with “medium quality” but the model’s performance had already been comparable to the best Chemprop model that cost 3,201 seconds for training. The best AutoGluon model that required the most computational resources only took 17 seconds to perform prediction. In comparison, the best Chemprop model cost 121 seconds for prediction.

#### Cytotoxicity

It is an unwanted property for antibiotic discovery, and it has become a key factor leading to the failed clinical trials of antibiotics^38^. Considering that some toxicity-related benchmarking sets are not only obsolescent for practical use but also has limited correlation to pharmacological toxicology^39^, we chose to retrieve training data from a recently published platform named DeepCancerMap^40^. It contains a subset comprised of 10,861 binary inhibitory labels on 32 normal cell lines from 9,668 compounds. With the same dataset, it become natural to compare model performance between AutoGluon and FP-GNN used in DeepCancerMap. Table S6 shows that for AutoGluon model training, RDKit 2D descriptor was slightly superior to Morgan2 fingerprint, with the MCC value of 0.754 versus 0.752. The best AutoGluon model significantly outperformed non-ensembled FP-GNN for which the MCC was only 0.536. However, we noticed that the AUC value of FP-GNN was greater than the highest AUC value achieved by AutoGluon, with the value of 0.914 versus 0.866.

#### Plasma Protein Binding

It is widely acknowledged that for small-molecule antibiotics, only free fraction of molecules can exert antibacterial effect, thus PPB can affect *in vivo* efficacy of antibiotics^41^. Therefore, accurate prediction of PPB can improve efficiency of screening candidates with appropriate binding affinity for plasma protein. In this case, a curated PPB dataset with 3,921 data points was retrieved from a recent study that reported IDL-PPBopt^42^, a deep learning model with graph attention mechanism for regressive prediction on PPB fraction value of compounds. AutoGluon was trained on this dataset and compared with IDL-PPBopt. Table S7 shows that RDKit 2D-based AutoGluon model achieved the best performance, with the MAE value of 0.062, RMSE value of 0.105 and *R*^2^ value of 0.861, which was superior to IDL-PPBopt model for which MAE was 0.075, RMSE was 0.112 and *R*^2^ was 0.841. However, it should be noted that ensemble learning was basically included for model training in AutoGluon while IDL-PPBopt was not ensembled.

#### Practical Application of Automated Molecular Design to Small-molecule Antibiotics

This section aims at simulating the application scenario of two tools in real-world drug discovery. 10,000 molecules sampled from the finetuned model were protonated between pH 7.3 and pH 7.5 to give 36,364 data points, which constituted the virtual library. The best configuration for each molecular property prediction model was adopted to train a prospective model with the whole dataset. Table S8 shows the performance of each model on the validation set. Then the models were applied to screening the virtual library. Figure S7 displays 9 potential antibacterial compounds for *E. coli.* with high probability of being active, and their cytotoxicity probability and PPB fraction are also annotated. All these compounds are not indexed in CAS SciFinder^43^ (accessed September 2023). 9,598 molecules with their predicted values are listed in Table S9, among which 42 molecules were predicted active (activity probability > 0.5), non-toxic (cytotoxicity probability < 0.5) and having appropriate PPB (predicted PPB fraction < 0.9), thus represented promising small-molecule antibiotics. Additionally, all the above predictions were performed on various device configurations to compare the runtime. Table S10 shows that even the most time-consuming task, i.e., PPB prediction, only cost about half an hour on an Apple computer, indicating that it is feasible to leverage these models for large-scale libraries on personal computer. Notably, the exploitation of GPU may not significantly accelerate the prediction. We found that most of the models trained with AutoGluon were CPU-intensive. To facilitate visualization of molecules, we designed a pop-up interface named “Molecular Visualization” (cf. Figure S8), where users can save either high-resolution bitmap or structure file of molecules.

## IMPLEMENTATION

AutoMolDesigner is an open-source software written in Python3. Freely accessible Qt Designer software was used to develop its GUI. The functionalities were realized by invoking application programming interfaces of PySide6. Another Python library named PyInstaller (version 5.12.0) was used to pack executable versions for both MacOS and Windows platform. The RNN based on PyTorch (version 1.13.1) was used for language modeling to implement deep molecular generation. The AutoML framework named AutoGluon (version 0.8.2) was adopted for molecular property prediction. Scikit-learn Python package (version 1.2.2) was used to calculate metrics for classification and regression^44^. A Python package named Descriptastorus (version 2.6.1) was used to compute all the molecular descriptors. RDKit^45^ Python package (version 2023.3.3) was used to perform basic molecular operations and visualization while MolVS (version 0.1.1) and Dimorphite-DL (version 1.3.2) was used for salt removal, molecular standardization, charge neutralization or protonation at a certain pH range.

## CONCLUSIONS

Combating antibiotic resistance somewhat relies on the discovery of small-molecule drugs with novel chemotypes. However, few user-friendly tools are publicly available for molecular design of small-molecule antibiotics. In this study, we presented an easy-to-use software dedicated to the AI-driven automated design of novel small-molecule antibiotics. We proposed an automated workflow that combined deep molecular generation with AutoML-based molecular property prediction. The benchmarking studies have proved that our tool is either superior or comparable to the other established models. The practical application has further shown that this software can produce novel molecules with high probability of being active, low probability of being toxic and appropriate PPB fraction. Notably, our software not only focuses on designing small molecules with whole cell-based antibacterial activity but also takes high antibiotic-likeness into consideration. In contrast, target-based computational methods, which leverage either experimentally-determined or computationally-modeled structures, have limited applicability in antibiotic discovery^46^. Ease of use is the other main feature of our software. It is equipped with an intuitive GUI that has been tested on mainstream operating systems. We have shown that users can have acceptable experience by using personal computers with limited computational capacity. Moreover, the curated datasets, the parameters of those trained models and user manual are made publicly available. We expect its wide application in novel small-molecule antibiotic discovery.

Currently, we have established the workflow for designing and screening small-molecule antibiotics. According to the aforementioned demonstration, this workflow could also be applied to other ESKAPE pathogens and other antibiotic-like properties. Our future work will be expanding the curated datasets and trained models for small-molecule antibiotics discovery and developing more useful modules of the software.

## ASSOCIATED CONTENT

### Data and Software Availability

The CLI version along with its source code is available at the open-source GitHub repository (https://github.com/taoshen99/AutoMolDesigner). The packaged GUI versions with user manual is free of charge at Zenodo (https://zenodo.org/record/8366085), where all the parameters of the trained models and the curated datasets that can be used for prospective study are also provided.

### Supporting Information

Materials and Methods, Software interface figures, data visualization figures, benchmarking results tables (Supporting_Information.docx); datasets made in model training and prediction results (Supplementary_Table.xlsx).

## AUTHOR INFORMATION

### Author Contributions

J.X. led the project, with the support from D.W. and S.W.. T.S. developed the source code and GUI. J.G. reviewed the code and made improvements. T.S., J.G. and Z.H. performed data collection and curation, and built ML models. T.S. and J.X. jointly performed the data analysis. J.G., Z.H., G.Z., Q.L. and X.S. tested the software and provided invaluable feedbacks. T.S. and J.X. wrote the manuscript. All authors reviewed the final version of the manuscript and provided important revisions.

### Notes

The authors declare no competing financial interest.

## Acknowledgements

This study was funded by the CAMS Innovation Fund for Medical Sciences (Grant No. 2021-I2M-1-069), the National Science and Technology Major Projects for Major New Drugs Innovation and Development (Grant No. 2018ZX09711001-012-003) and the Program for Foreign Talent of Ministry of Science and Technology of the People’s Republic of China (Grant No. G2021194015L).

## Supporting Information

**Figure S1.**
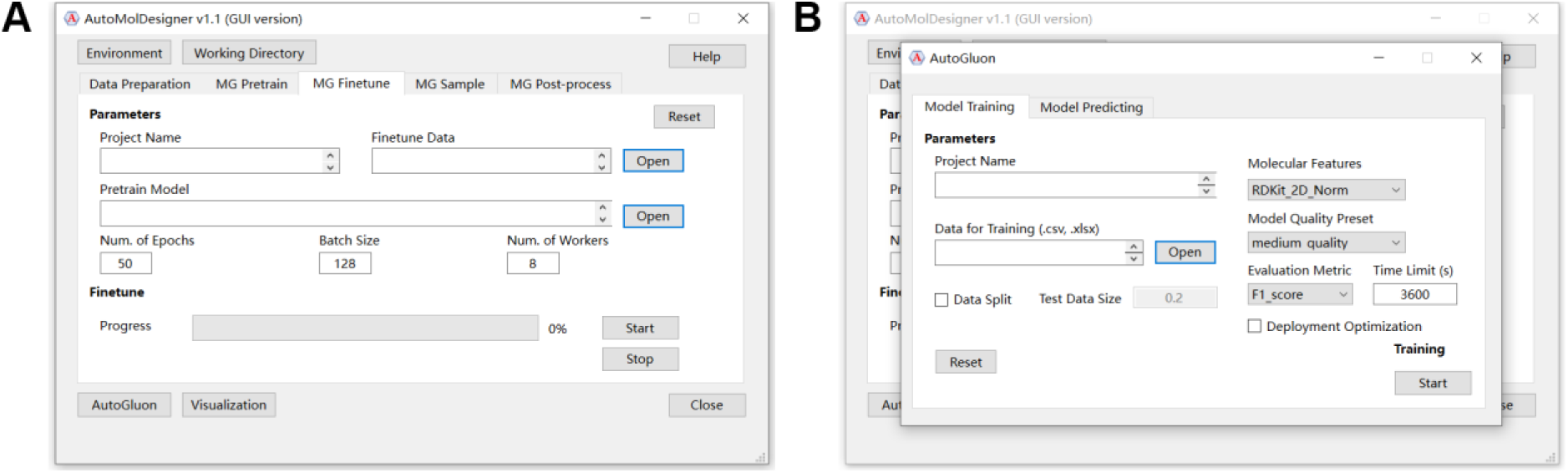
Two functional windows of AutoMolDesigner. (**A**) Main window: deep molecular generation for *de novo* drug design; (**B**) Pop-up window: automated machine learning for molecular property prediction.

**Figure S2.**
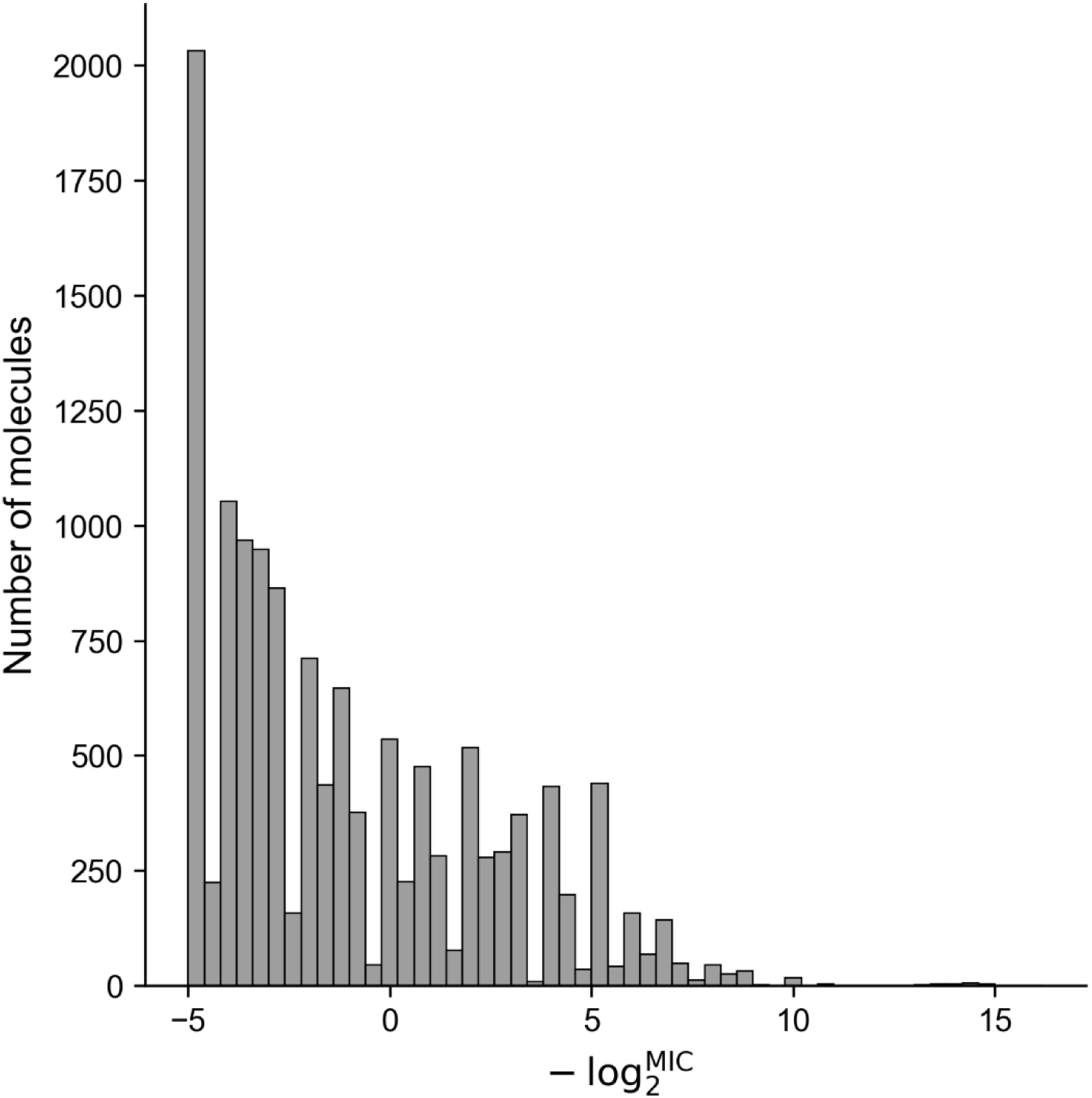
Distribution of antibacterial activity (minimum inhibitory concentration, MIC, μg/mL). The value was transformed as -log_2_^MIC^.

**Figure S3.**
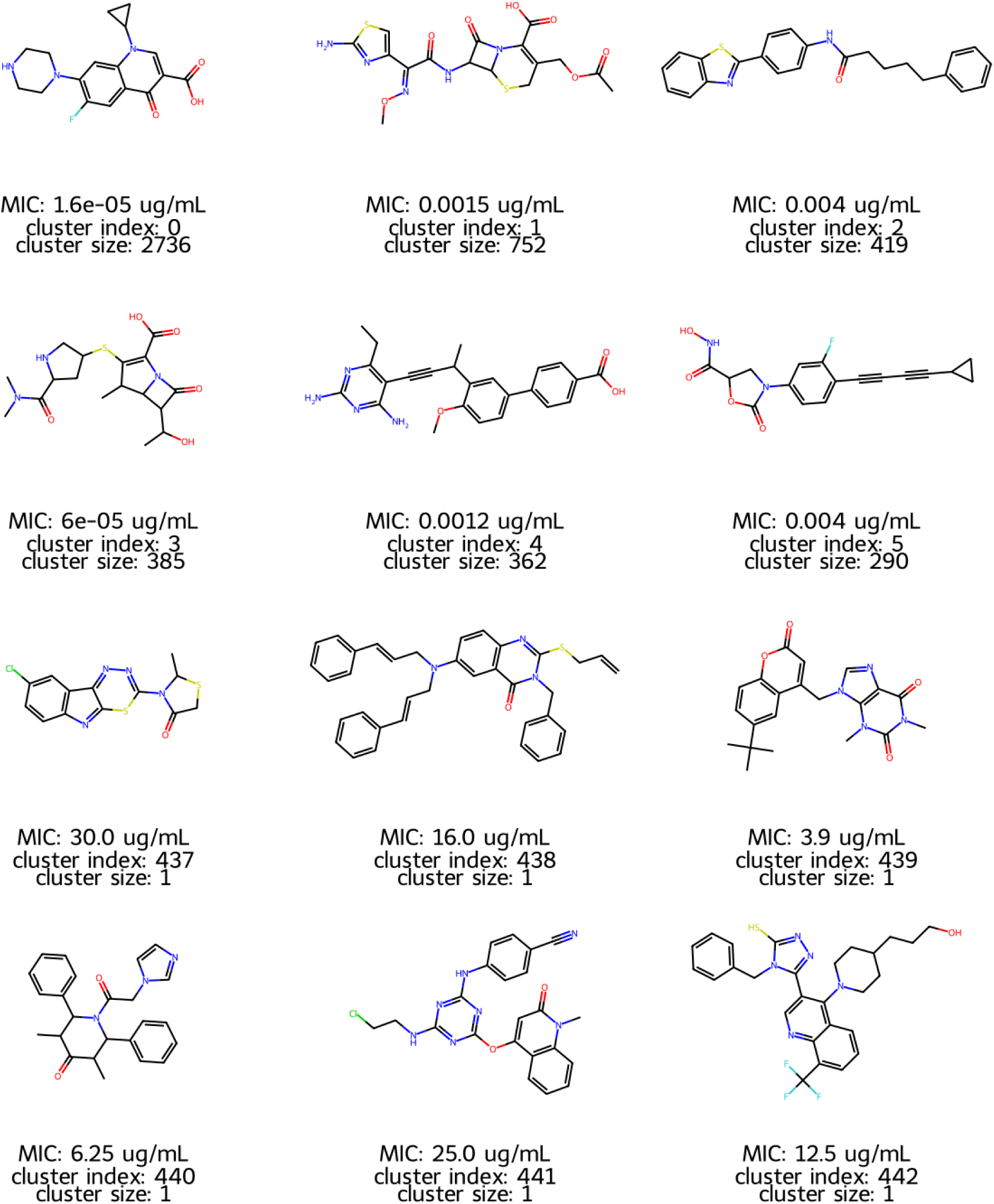
Compounds annotated with MIC, cluster index and cluster size from 12 selected clusters. The cluster index can be referred to Table S2.

**Figure S4.**
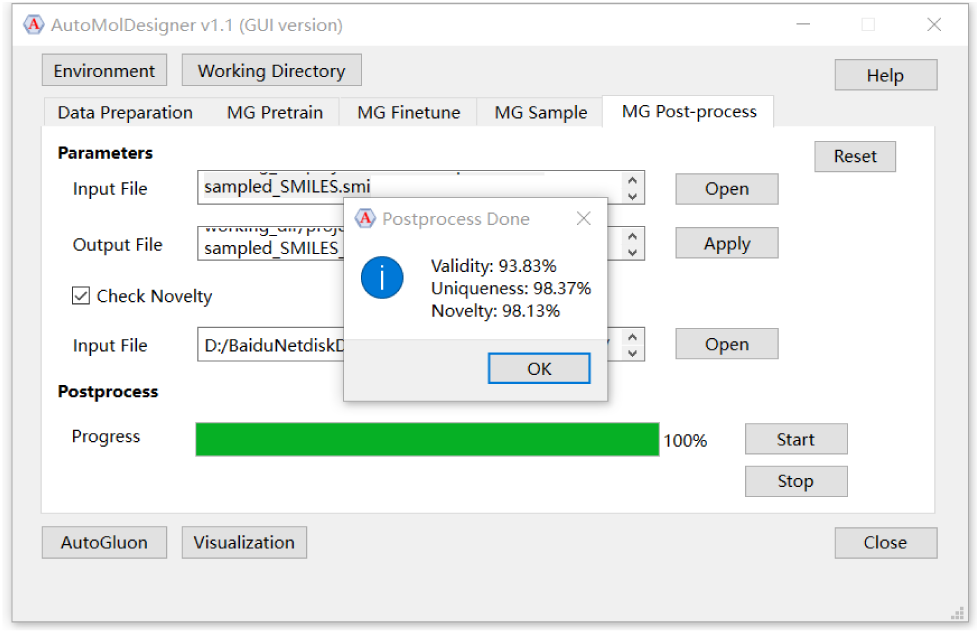
“Validity”, “Uniqueness” and “Novelty” of 1,000 molecules sampled from the finetuned model at “MG Post-process” tab.

**Figure S5.**
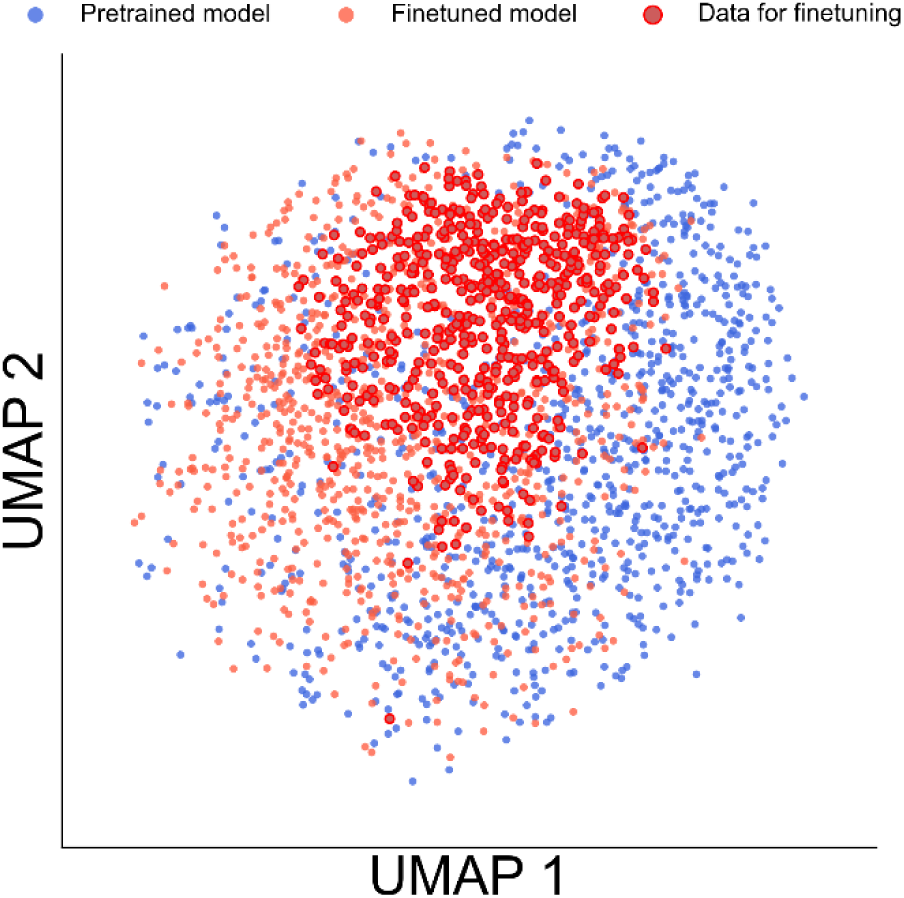
UMAP visualization of molecules embedded in chemical space characterized by FCFP_6. 967 molecules sampled from the pretrained model are colored blue, 910 molecules sampled from the finetuned model are colored orange and 570 molecules from “Datasets for Finetuning” are colored dark red.

**Figure S6.**
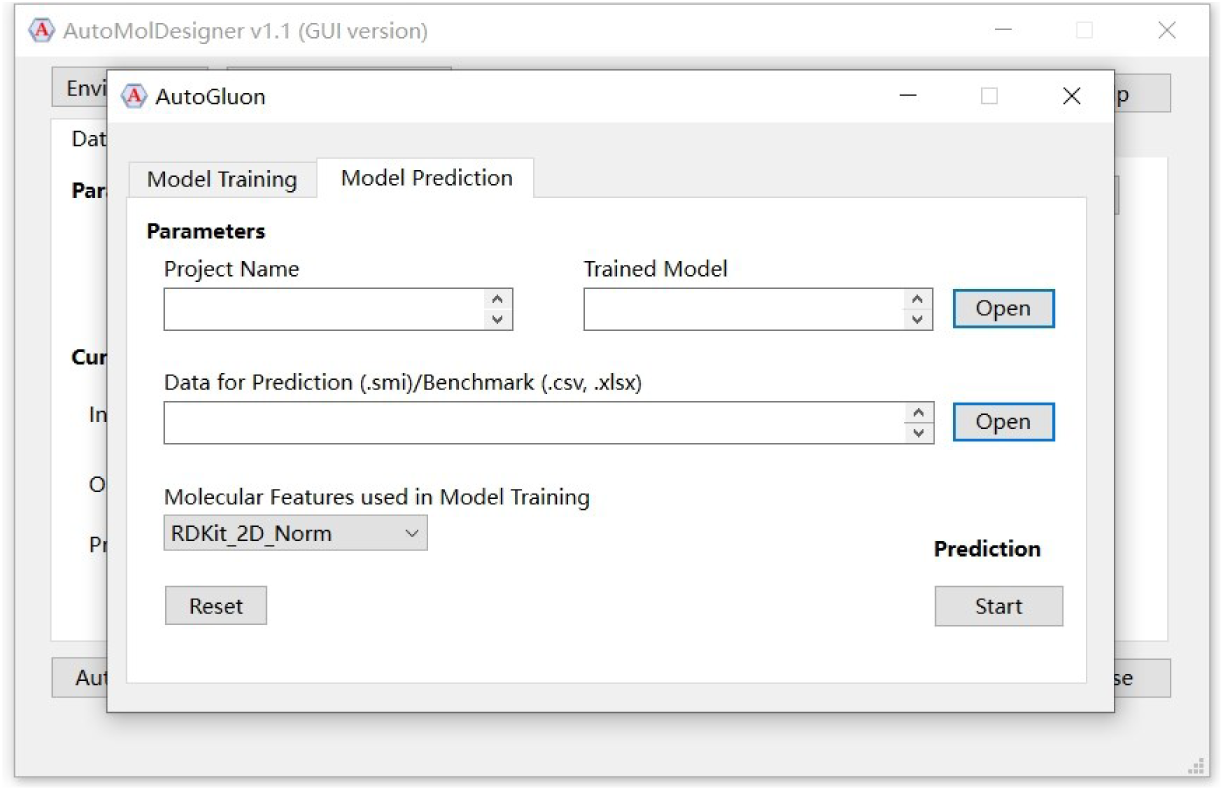
“Model Prediction” interface of the AutoGluon module.

**Figure S7.**
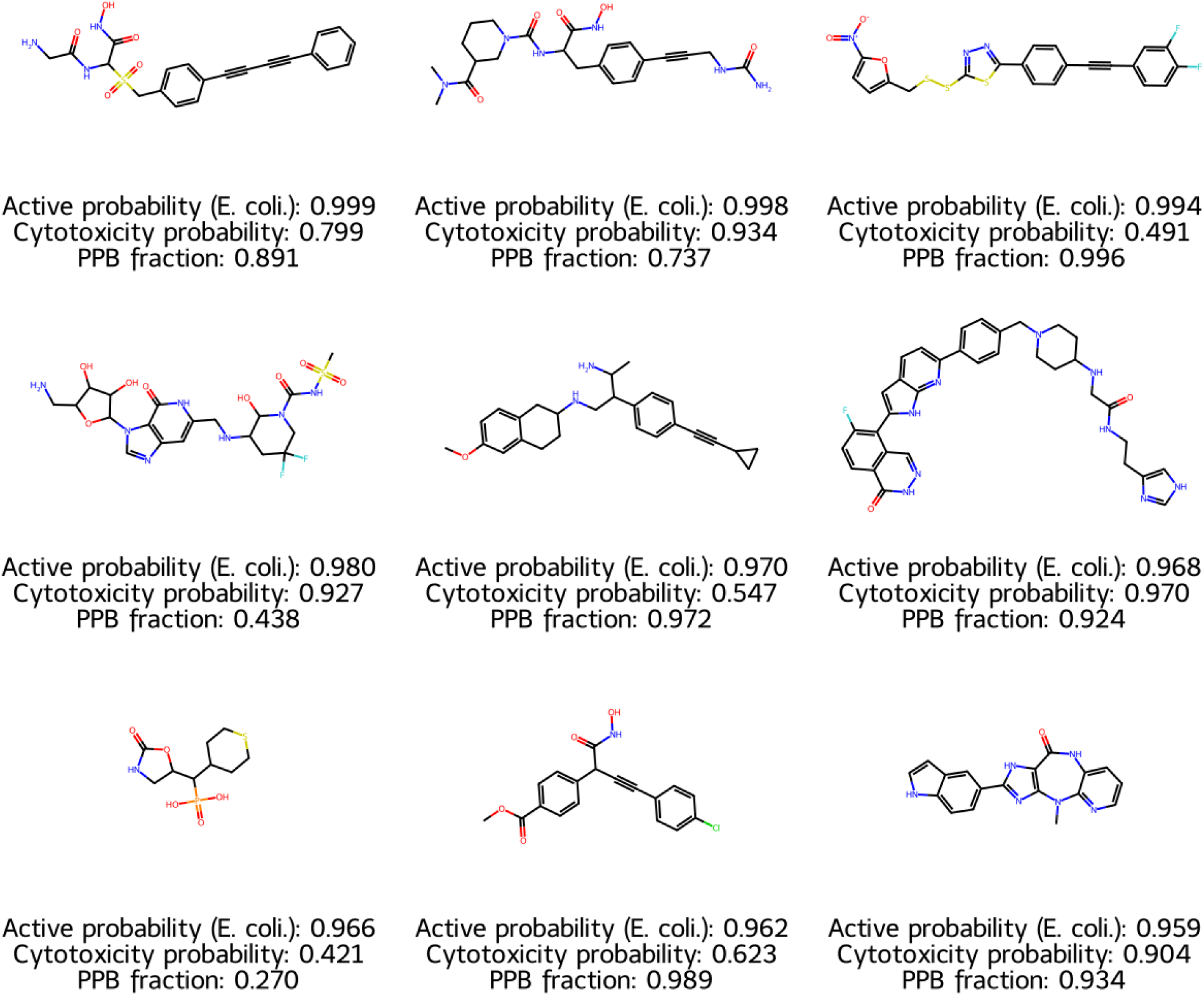
9 designed antibacterial compounds with high active probability, alongside the cytotoxicity probability and PPB fraction.

**Figure S8.**
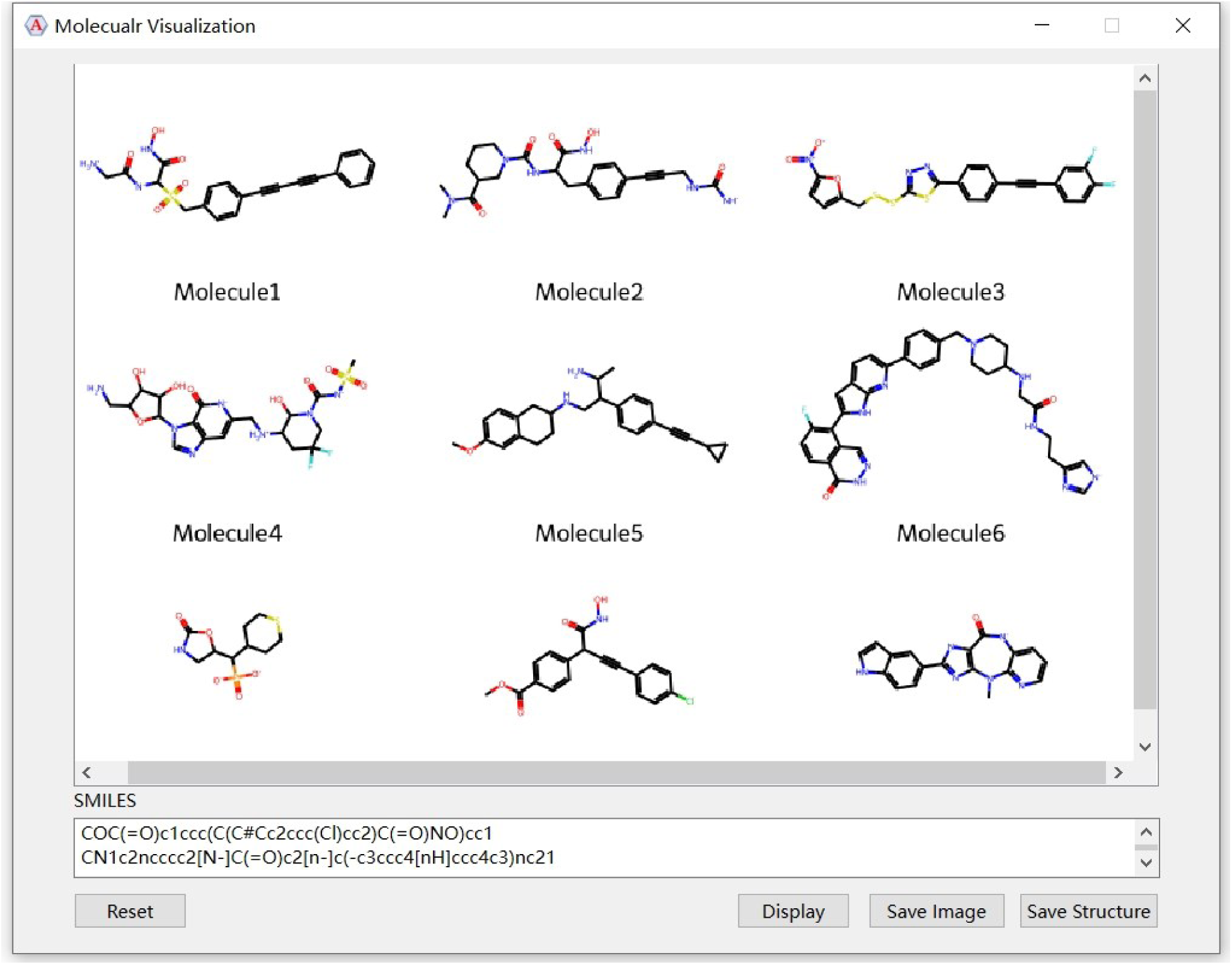
“Molecular Visualization” interface popping up from the main interface.

**Table S1.**
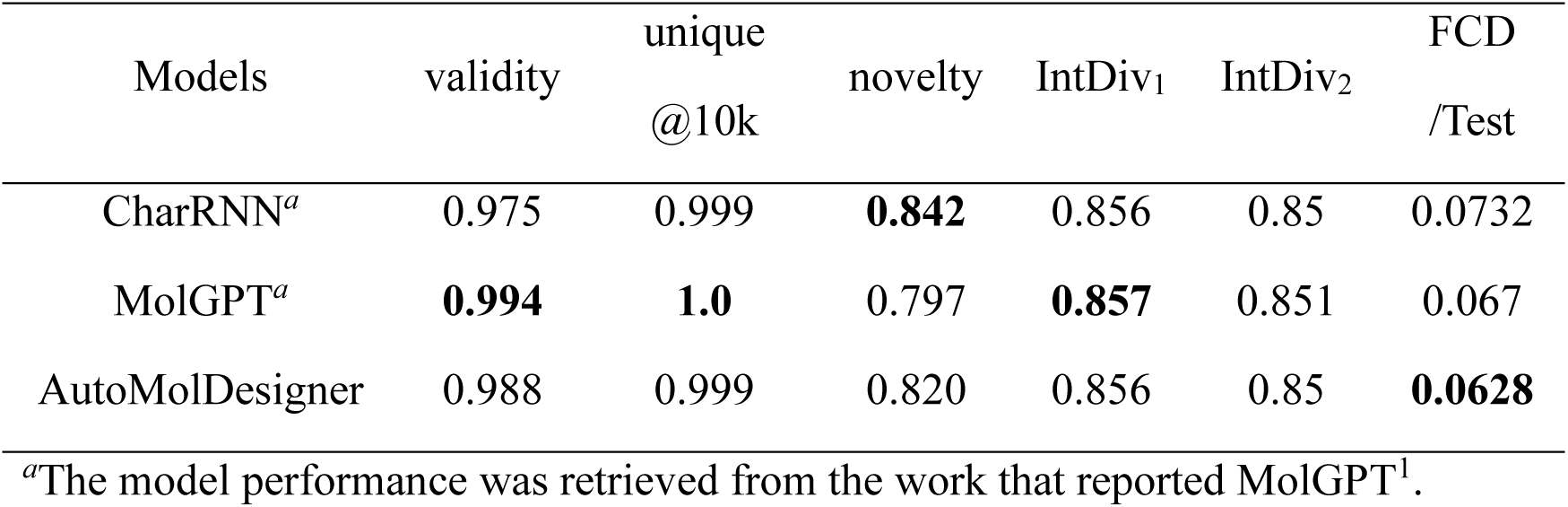
Performance comparison between our remastered RNN from AutoMolDesigner and other deep molecular generation models, i.e., CharRNN and MolGPT, with MOSES as the benchmark. The best value for each metric is highlighted in bold.

**Table S3.**
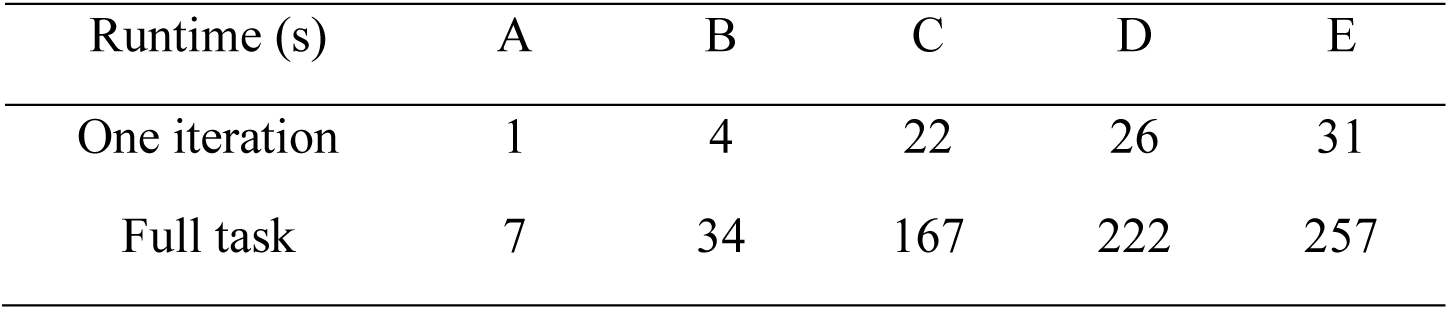
Runtime of model finetuning with various device configurations. Configuration (**A**) Ubuntu server: CPU [Intel(R) Core(TM) i9-12900K @ 3.20GHz] with GPU [NVIDIA GA102 (GeForce RTX 3090 24GB)]; (**B**) Ubuntu server: CPU [Intel(R) Core(TM) i9-12900K @ 3.20GHz]; (**C**) Windows laptop: CPU [AMD Ryzen(TM) 7 5800H @ 3.20GHz] with GPU [NVIDIA GA106 (GeForce RTX 3050Ti Mobile 4GB)]; (**D**) Windows laptop: CPU [AMD Ryzen(TM) 7 5800H @ 3.20GHz]; (**E**) Apple Mac mini: CPU (Apple M2 @ 3.49GHz).

| Runtime (s) | A | B | C | D | E |
| --- | --- | --- | --- | --- | --- |
| One iteration | 1 | 4 | 22 | 26 | 31 |
| Full task | 7 | 34 | 167 | 222 | 257 |

**Table S5.** Antibacterial activity prediction – comparative study between Chemprop and AutoGluon. The best value for each metric is highlighted in bold.

| Models |  | ACC<sup>a</sup> | AUC<sup>b</sup> | MCC<sup>c</sup> | F1 score | Training time<sup>g</sup> (s) | Predicting time (s) |
| --- | --- | --- | --- | --- | --- | --- | --- |
| **Chemprop** |  |  |  |  |  |  |  |
| baseline |  | 0.974 | 0.852 | 0.443 | 0.456 | 155 | 4 |
| baseline ensemble<sup>d</sup> |  | 0.979 | 0.873 | 0.524 | 0.533 | 2782 | 48 |
| RDKit 2D feature<sup>e</sup> |  | 0.980 | 0.864 | 0.520 | 0.521 | 457 | 79 |
| RDKit 2D feature ensemble |  | 0.981 | 0.877 | 0.552 | 0.557 | 3201 | 121 |
| Morgan2 feature |  | 0.978 | 0.903 | 0.506 | 0.511 | 161 | 5 |
| Morgan2 feature ensemble |  | 0.979 | 0.914 | 0.532 | 0.540 | 2925 | 52 |
| **AutoGluon** |  |  |  |  |  |  |  |
| RDKit_2D<sup>f</sup> | baseline | 0.983 | 0.879 | 0.572 | 0.555 | **27** | **0.08** |
|  | best | 0.981 | 0.930 | 0.567 | 0.573 | 1687 | 10 |
| Morgan 2 | baseline | 0.982 | 0.870 | 0.544 | 0.520 | 67 | 0.9 |
|  | best | **0.987** | **0.937** | **0.687** | **0.671** | 3365 | 17 |

<sup>a</sup>ACC: Accuracy. <sup>b</sup>AUC: Area under receiver operating characteristic. <sup>c</sup>MCC: Matthews correlation coefficient. <sup>d</sup>The ensemble consisted of 20 independently trained models. <sup>e</sup>Chemprop was enhanced by molecular-level features that are additional molecular descriptors. <sup>f</sup>This molecular descriptor named normalized RDKit\_2D was the same as that used for Chemprop enhancement. <sup>g</sup>For Chemprop, the training time did not include the time for hyperparameters optimization.

**Table S6.** Cytotoxicity prediction – comparative study between FP-GNN and AutoGluon. This benchmark contained 32 tasks, and their results were averaged for comparison. The best value for each metric is highlighted in bold.

| Models |  | Sensitivity | Specificity | ACC<sup>a</sup> | AUC<sup>b</sup> | MCC<sup>c</sup> |
| --- | --- | --- | --- | --- | --- | --- |
| **FP-GNN** |  |  |  |  |  |  |
| Reported model |  | 0.813 | 0.423 | 0.722 | **0.914** | 0.536 |
| **AutoGluon** |  |  |  |  |  |  |
| RDKit_2D<sup>d</sup> | baseline | 0.952 | 0.643 | 0.853 | 0.818 | **0.754** |
|  | best | 0.953 | **0.662** | **0.877** | 0.866 | 0.750 |
| Morgan2 | baseline | 0.948 | 0.593 | 0.837 | 0.819 | 0.723 |
|  | best | **0.962** | 0.632 | 0.872 | 0.864 | 0.752 |

<sup>a</sup>ACC: Accuracy. <sup>b</sup>AUC: Area under receiver operating characteristic. <sup>c</sup>MCC: Matthews correlation coefficient. <sup>d</sup>Normalized version.

**Table S7.** Plasma protein binding prediction – comparative study between IDL-PPBopt and AutoGluon. The best value for each metric is highlighted in bold.

| Models | | MAE<sup>a</sup> | RMSE<sup>b</sup> | $R^2$<sup>c</sup> | Training time (s) | Predicting time (s) |
| --- | --- | --- | --- | --- | --- | --- |
| **IDL-PPBopt** |  |  |  |  |  |  |
| Reported model |  | 0.075 | 0.112 | 0.841 | -<sup>e</sup> | 2 |
| **AutoGluon** |  |  |  |  |  |  |
| RDKit_2D<sup>d</sup> | baseline | 0.068 | 0.110 | 0.846 | **25** | **0.7** |
|  | best | **0.062** | **0.105** | **0.861** | 3511 | 8 |
| Morgan2 | baseline | 0.084 | 0.134 | 0.772 | 146 | 1.0 |
|  | best | 0.074 | 0.124 | 0.806 | 3542 | 15 |

<sup>a</sup>MAE: Mean absolute error. <sup>b</sup>RMSE: root mean squared error. <sup>c</sup> $R^2$ : $R$ square. <sup>d</sup>Normalized version.
<sup>e</sup>The official script for training is not available in the published work.

**Table S8.** Validation set performance of three molecular property prediction models applied to screening the virtual library.

| Metric<sup>a</sup> | Antibacterial activity | Cytotoxicity | Plasma protein binding |
| --- | --- | --- | --- |
| F1 score | 0.645 | 0.894 | _<sup>b</sup> |
| MAE | _<sup>b</sup> | _<sup>b</sup> | 0.062 |

<sup>a</sup>The “Evaluation Metric” argument for AutoGluon model training. <sup>b</sup>Not applied.

**Table S10.** Runtime of predicting 36,364 data points on three tasks with various device configurations (see definitions in Table S3).

| Runtime (s) | A | B | C | D | E |
| --- | --- | --- | --- | --- | --- |
| Antibacterial activity | 77 | 80 | 252 | 254 | 106 |
| Cytotoxicity | 167 | 170 | 300 | 313 | 211 |
| Plasma protein binding | 666 | 660 | 1006 | 1029 | 1744 |

### MATERIALS AND METHODS

#### Data Collection and Curation

In the section of “Tool I: Generating Focused Library for Small-molecule Antibiotics”, all the training data were retrieved from ChEMBL^2^ (version 32, https://www.ebi.ac.uk/chembl/, accessed March 2023). For pretraining, 764,373 small molecules (SMILES) whose annotated bioactivities (“K_d_”, “K_i_”, “K_b_”, “IC50”, “EC50”) were no higher than 10 μM were obtained^3^. After the standard curation including de-duplication, salts removal and molecular standardization, they were further augmented by 10 folds to give 7,230,290 data points (“Dataset for Pretraining”). For finetuning, 15,795 small molecules whose annotated minimum inhibitory concentration (MIC) values were no higher than 32 μg/mL for *Escherichia coli* (*E. coli*) were obtained after “NaN” dropping and de-duplication. Apart from the standard curation, molecules whose molecular weight were greater than 600 or number of rotatable bonds were more than 20^4, 5^ were also removed to give 13,243 data points. Structural clustering based on Morgan2 fingerprints using Butina algorithm with 0.75 (Tanimoto similarity) as the threshold was carried out to give 570 clusters^6^. The most active molecule from each cluster was collected to constitute the final finetuning datasets (“Dataset for Finetuning”).

In the section of “Tool II: AutoML-based Molecular Property Prediction for Small-molecule Antibiotics”, MUBD-DecoyMaker2.0^7^ was used to make the dataset used for training and predicting antibacterial activity against *E. coli.*. Briefly, molecules labeled with “high activity” (MIC ≤ 1 μg/mL) from 13,243 data points were protonated at pH ranging from 7.3 to7.5 by Discovery Studio^8^ (version 2016) to give 5,021 data points, which served as the input for MUBD-DecoyMaker2.0. The final MUBD dataset was comprised of 501 unbiased ligands and 19,539 unbiased decoys. The dataset used for cytotoxicity prediction was retrieved from a subset in DeepCancerMap^9^. This server contains 32 subsets comprised of compounds with annotated binary toxicity labels against normal cell lines. We followed its train-test splitting way to train 32 individual models. The dataset used for plasma protein binding (PPB) prediction was basically retrieved from IDL-PPBopt^10^, which contains 3,921 records. Its published splits of training, validating and testing sets were strictly followed.

There were two sets of molecules generated by the pretrained and finetuned model, respectively. One set contained 1,000 data points that were used for chemical space visualization. The other set of molecules including 10,000 data points was generated by the finetuned model for the simulated experiment of virtual screening. All the post-processing procedures implemented in “MG Post-process” tab were performed on these sets of molecules to remove invalid and duplicate molecules. Moreover, the recalled molecules in “Dataset for Finetuning” were also removed to ensure novelty. The data points for the simulated experiment of virtual screening were also protonated at pH ranging between 7.3 and 7.5. Eventually, 36,364 data points were generated.

#### Deep Molecular Generation

The deep molecular generation implemented in our software program belonged to the well-established chemical language model^11, 12^. Briefly, SMILES sequences were firstly tokenized to atomic symbols that constituted a vocabulary together with tokens for starting, ending and padding. Next, an embedding layer of 128 dimensions followed by three-layer long-short term memory (LSTM) with 512 units were used to process these tensors, and a final dense layer with the same dimension as the length of the vocabulary was used to put out un-softmaxed value for each token. Dropout and layer normalization and gradient clipping were used to prevent overfitting. The model was trained in an auto-regressive manner with cross entropy loss, default Adam optimizer in PyToch and learning rate schedule of cosine annealing (T_max = 64 for pretraining and T_max = 32 for finetuning). For pretraining, the dataset was randomly split into a training set and a validation set with the ratio of 99:1 while for finetuning, the ratio was 4:1. The batch size for pretraining (512) was also larger than that for finetuning (128). Moreover, the parameters of the embedding layer learned from model pretraining were frozen during backward propagation in model finetuning. For model sampling, the SMILES sequence was generated token by token until the ending token occurred or the maximum token length of 128 was reached. A widely used technique named “Temperature sampling”^11^ was also included to control molecular novelty by stretching the probability distribution of candidate tokens.

#### Molecular Property Prediction

The molecular descriptors and other optional arguments for AutoGluon^13^ model training and prediction are illustrated in the main text. Herein, we describe the detailed settings of retrospective studies for three molecular property prediction models. In all these studies, AutoGluon models adopted normalized RDKit 2D descriptors and Morgan2 fingerprints. Two training modes were applied, i.e., “baseline” mode denoting that model was trained in the default configuration provided by AutoGluon, and “best” mode denoting that model was trained in the configuration of “Time limit (s)” = 3600 and “Model Quality Presets” = “best model quality”. All evaluation metrics were calculated by Scikit-learn^14^ Python package (version 1.2.2). For antibacterial activity prediction, hyperparameter optimization was firstly performed for Chemprop^15^ with 80% molecules from the MUBD dataset as the training set and through five-fold cross-validation with 10 iterations. Afterwards, the test set (20% remaining molecules) was used to compare the performances of optimized Chemprop and AutoGluon models. For cytotoxicity prediction, the average performance of FP-GNN on 32 tasks were used for comparison, and the AutoGluon models were benchmarked in the same manner. For PPB fraction prediction, the performance of IDL-PPBopt^10^ was retrieved from the published results while AutoGluon was benchmarked with the training and test sets from IDL-PPBopt. All the benchmarking experiments were conducted in triplicate. It is worth noting that the last three prospective models trained with the full datasets were optimized by “Deployment Optimization”.

**Figure S9.**
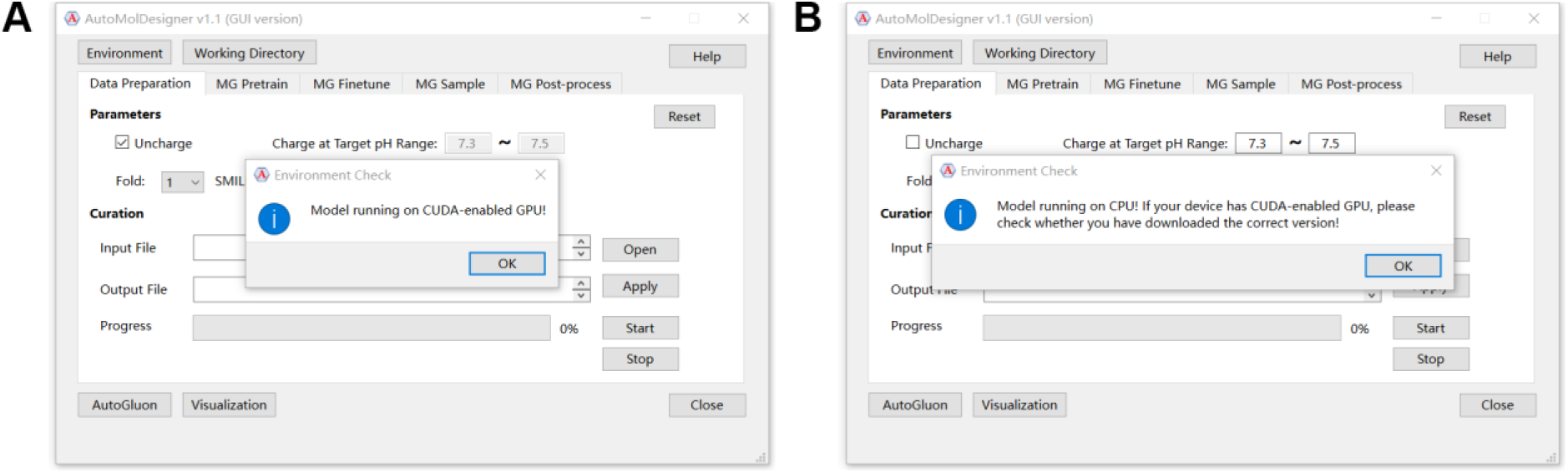
Detect running environment. (**A**) Model runs on CUDA-enabled GPU. (**B**) Model runs on CPU.

#### Other Useful Modules

The usage and implementation of other useful modules including data curation, model benchmarking and molecular visualization are described below. To begin with, there is a button located at the upper left corner of the “Environment” interface. It can be clicked to check whether the application is central processing unit (CPU) version or graphical processing unit (GPU) version. This functionality was realized by the PyTorch function in checking availability of compute unified device architecture (CUDA). The model will be run on CPU if CUDA-enabled GPU is not available (Figure S9). The second button called “Working Directory” can be used to set the current working directory. As a result, all the data generated will be saved under the directory of “your working directory/projects/your project name”. It should be noted that the “Reset” button is available in each tab for resetting. In terms of “Data preparation” tab, several class methods of MolVS (version 0.1.1) and Dimorphite-DL^16^ (version 1.3.2) including “LargestFragmentChooser().choose()”, “Uncharger().uncharge()”, “Standardizer().standardize()”, and “DimorphiteDL().protonate()” were used to implement salts removal, charge neutralization, standard molecular standardization and protonation at a certain pH range. Moreover, “SmilesEnumerator().randomize_smiles()” that is a class method implemented in a standalone Python script called “SmilesEnumerator.py” was used to obtain the single randomized SMILES of a molecule. For “MG Post-process” tab, all the curation is based on Python built-in functions for data manipulation. “Molecular Visualization” tab is another supplementary module that provides a convenient way to show the designed molecules, and it was implemented by the “rdkit.Chem.Draw.MolsToGridImage()” function. Additionally, all the other molecular manipulations such as basic sanitization and molecular format conversion between SMILES and Structural Data were all made by RDKit^17^ (version 2023.3.3). With its built-in dataframe, Pandas (version 1.5.3) was used to process tabular data in the form of Comma-Separated Values (*.csv) or Microsoft Excel (*.xlsx).

#### Graphical User Interface Implementation

As mentioned in the section of “Implementation”, the graphical user interface (GUI) of AutoMolDesigner was designed based on Qt platform wrapped by PySide6 (version 6.5.1.1). The GUI code, the functional source code, and the related packages were packed into an executable and distributable standalone software by PyInstaller (version 5.12.0) for both MacOS and Windows operating systems (for Linux users, GUI version is currently not available, as command-line interface version that basically has the same operating logics can serve as an alternative). In detail, each long-term task will run on an independent thread through “QThread” and “QThreadPool” classes of PySide6 to maintain responsiveness of the application. For molecular generation in the main interface, the progress bar is provided for each tab to indicate the progress of current task, while the adjacent “Stop” button can be used for manual termination. But it should be noted that pretraining and finetuning tasks will not stop immediately after the click of this button and will stop only when the current iteration is completed. For molecular property prediction in the “AutoGluon” interface, a logging dialogue will pop up during model training or prediction. If users try to close this dialogue before the task is completed, a message box will pop up as a reminder. Only if further confirmation is made according to the message box, the task can be manually terminated (Figure S10).

**Figure S10.**
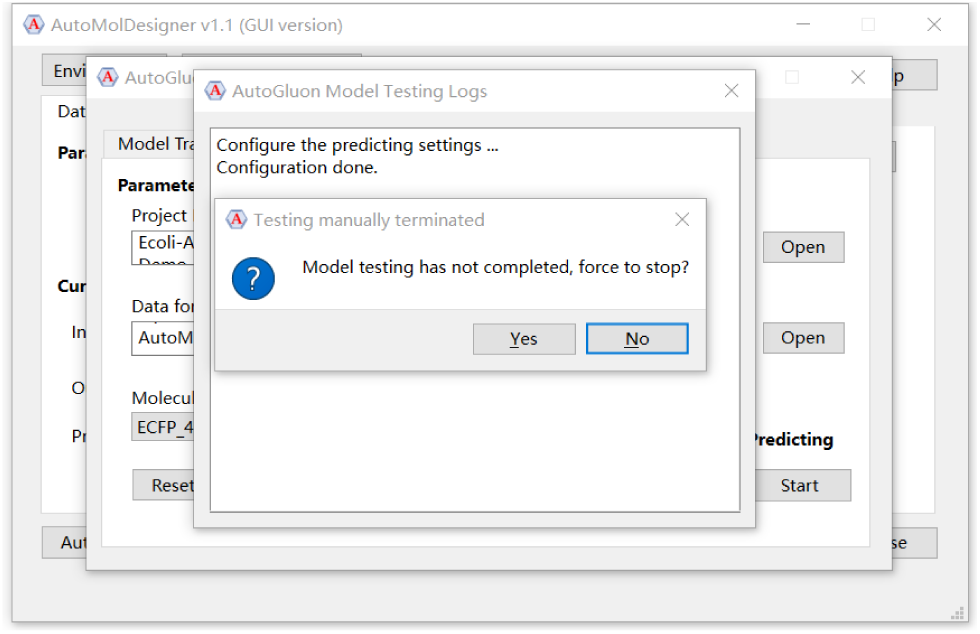
The message box pops up from the “AutoGluon” interface when users try to manually terminate the task.

